## Supplementary material for "Signals from the head and germinative region differentially regulate regeneration competence of the tapeworm *Hymenolepis diminuta*": Table S1

**Table S1. *βcatenin* paralogs in *H. diminuta.***

| ***βcat1*** | | | | |
| --- | --- | --- | --- | --- |
| Query sp. | Query Accession | Reference | Best match to *H. diminuta** | E-value |
| *E. multilocularis* | EmuJ_001007700 | Montagne et al., 2019 | WMSIL1_LOCUS14475 | 0 |
| *S. mansoni* | Smp_023550 | Montagne et al., 2019 | WMSIL1_LOCUS14475 | 0 |
| *S. mediterranea* | ABW79875.1 | Su et al., 2017 | WMSIL1_LOCUS14475 | 2.40E-98 |
| ***βcat2*** | | | | |
| Query sp. | Query Accession | Reference | Best match to *H. diminuta** | E-value |
| *E. multilocularis* | EmuJ_001103600 | Montagne et al., 2019 | WMSIL1_LOCUS880 | 7.20E-148 |
| *S. mansoni* | Smp_173990 | Montagne et al., 2019 | WMSIL1_LOCUS880 | 5.10E-90 |
| *S. mediterranea* | ABW79874.1 | Su et al., 2017 | WMSIL1_LOCUS880 | 2.80E-33 |
| ***βcat3/4*** | | | | |
| Query sp. | Query Accession | Reference | Best match to *H. diminuta** | E-value |
| *E. multilocularis* | EmuJ_000572500 | Montagne et al., 2019 | WMSIL1_LOCUS6332 | 0 |
| *S. mansoni* | Smp_134000 | Montagne et al., 2019 | WMSIL1_LOCUS6332 | 4.00E-18 |
| *S. mediterranea* | KY196224 | Su et al., 2017 | none |  |
| *S. mediterranea* | KY196225 | Su et al., 2017 | none |  |
| *Genome assembly PRJEB30942 | |  |  |  |

**Table S2. Primers and transcripts used in this study.**

(attached excel file)

**Table S3. Source data used in this study.**

(attached excel file)
